## Supplemental Material for "Automated detection and quantification of two-spotted spider mite life stages using computer vision for high-throughput *in vitro* assays"

### Supplemental materials

#### Supplemental model testing and optimization methods

##### Methods

To examine the effect of backbone architecture on detection accuracy and efficiency, we trained and evaluated multiple object detection models, including YOLOv8 (Nano and Large), YOLO11 (Nano and Large), RT-DETR-L, and Faster R-CNN with two backbone configurations [45,46]. All models were trained on the same v209 five-class dataset annotated in YOLO bounding-box format, using a fixed random seed to ensure reproducibility. Ultralytics models were trained using the native training pipeline with default data augmentation and optimization settings. Faster R-CNN models (ResNet-50-FPNv2 and MobileNetV3-FPN backbones, both from Torchvision) were initialized from ImageNet-pretrained weights and trained with standard Torchvision hyperparameters: stochastic gradient descent (SGD) with momentum 0.9, weight decay 0.0005, and a multistep learning rate schedule (initial learning rate = 0.005). All models were trained for 50 epochs at an input resolution of 1024 × 1024 pixels, using a batch size of two and two dataloader workers. Model performance was evaluated using mean average precision at 50% intersection-over-union (mAP50), mean average precision across IoU thresholds from 0.50 to 0.95 (mAP50–95), and, for efficiency, mean per-image inference latency converted to approximate frames per second (FPS).

To evaluate the effect of image resolution on detection performance, images from the v209 test dataset were down sampled to 819, 512, and 256 pixels on the long edge and then upscaled back to 1024 × 1024 pixels for inference using model v209. All datasets were evaluated under identical inference settings (confidence threshold = 0.5, seed = 42). mAP50, mAP50-95, precision and recall were compared across resolutions to assess performance degradation with reductions in spatial detail. Because detection of *Viable_egg* represents the smallest object scale, changes in its accuracy were used to illustrate resolution sensitivity: individual eggs measured approximately 14 × 14 pixels at 1024 px, 11 × 11 pixels at 819 px, 7 × 7 pixels at 512 px, and 3.5 × 3.5 pixels at 256 px.

##### Results and Discussion

Across models trained on the *v209* five-class dataset, larger YOLO variants (YOLOv8-L and YOLO11-L) consistently outperformed their smaller counterparts in both mAP50 and mAP50–95 (Table S3). YOLO11-L achieved the highest overall performance (mAP50 = 0.836, mAP50–95 = 0.535), while also maintaining competitive inference speed (50.9 FPS) relative to its parameter count (25.3 M). YOLOv8-L followed closely (mAP50 = 0.830, mAP50–95 = 0.510), showing similar accuracy but with a larger parameter footprint.

The transformer-based RT-DETR-L achieved moderate detection performance (mAP50 = 0.770, mAP50–95 = 0.486) but operated at a substantially lower inference speed (36.7 FPS), highlighting the trade-off between transformer precision and real-time performance. Among the smaller models, YOLOv8-N and YOLO11-N exhibited the highest efficiency, with YOLOv8-N achieving 109.8

FPS and YOLO11-N reaching 80.6 FPS, though at a modest cost in accuracy (mAP50–95 = 0.454 and 0.435, respectively).

The Faster R-CNN architectures showed markedly lower accuracy under identical training conditions. The ResNet-50-FPNv2 variant achieved mAP50 = 0.588 and mAP50–95 = 0.336, while the lighter MobileNetV3-FPN model performed poorly overall (mAP50 = 0.244, mAP50–95 = 0.098) despite its smaller size. These results underscore the substantial performance advantage of recent YOLO architectures, particularly YOLO11-L, for high-resolution multi-class detection tasks in this dataset. Consequently, YOLO11 was selected as the base architecture for subsequent model refinement and application experiments.

Detection accuracy remained largely stable following moderate reductions in resolution, with mAP50 declining only slightly from 0.84 at 1024 px to 0.81 at 819 px (see Tables S4-S5 below). However, substantial performance losses occurred below 50% of the native scale. At 512 px, mAP50 dropped to 0.72, and at 256 px it declined to 0.46, accompanied by a sharp decrease in recall (0.24). Class-level results showed that small-object classes such as *Viable_egg* were most affected, while larger forms (*Adult_female*) remained comparatively robust. Overall, YOLO11l tolerated modest reductions in spatial detail but exhibited a large decline in detection accuracy once object size approached the model's lower stride limit (8x8 pixels), suggesting a practical resolution threshold between approximately 512 and 819 pixels for reliable inference.

The resolution ablation results indicate that YOLO11-l maintained robust detection performance until image detail was reduced below approximately half of the training scale. At 819 × 819 pixels, overall mAP50 remained within 3% of the 1024-pixel baseline, but accuracy declined sharply at 512 pixels and collapsed at 256 pixels. This pattern reflects the relationship between object size and the model's spatial sampling capacity.

In the original images, a *Viable_egg* occupied roughly 13 × 13 pixels at 1024 resolution, 10 × 10 pixels at 819, and only about 7 × 7 pixels at 512. Because YOLO's smallest detection head operates on an 8-pixel stride, features smaller than ~10 pixels across are represented by fewer than two grid cells and become poorly encoded. Consequently, the smaller classes exhibited the steepest losses in recall as resolution decreased. This suggests that image acquisition should target a pixel resolution sufficient to represent the smallest objects of interest with at least 10–12 pixels across their major axis, corresponding to roughly 0.5 µm per pixel.

### Supplemental figures

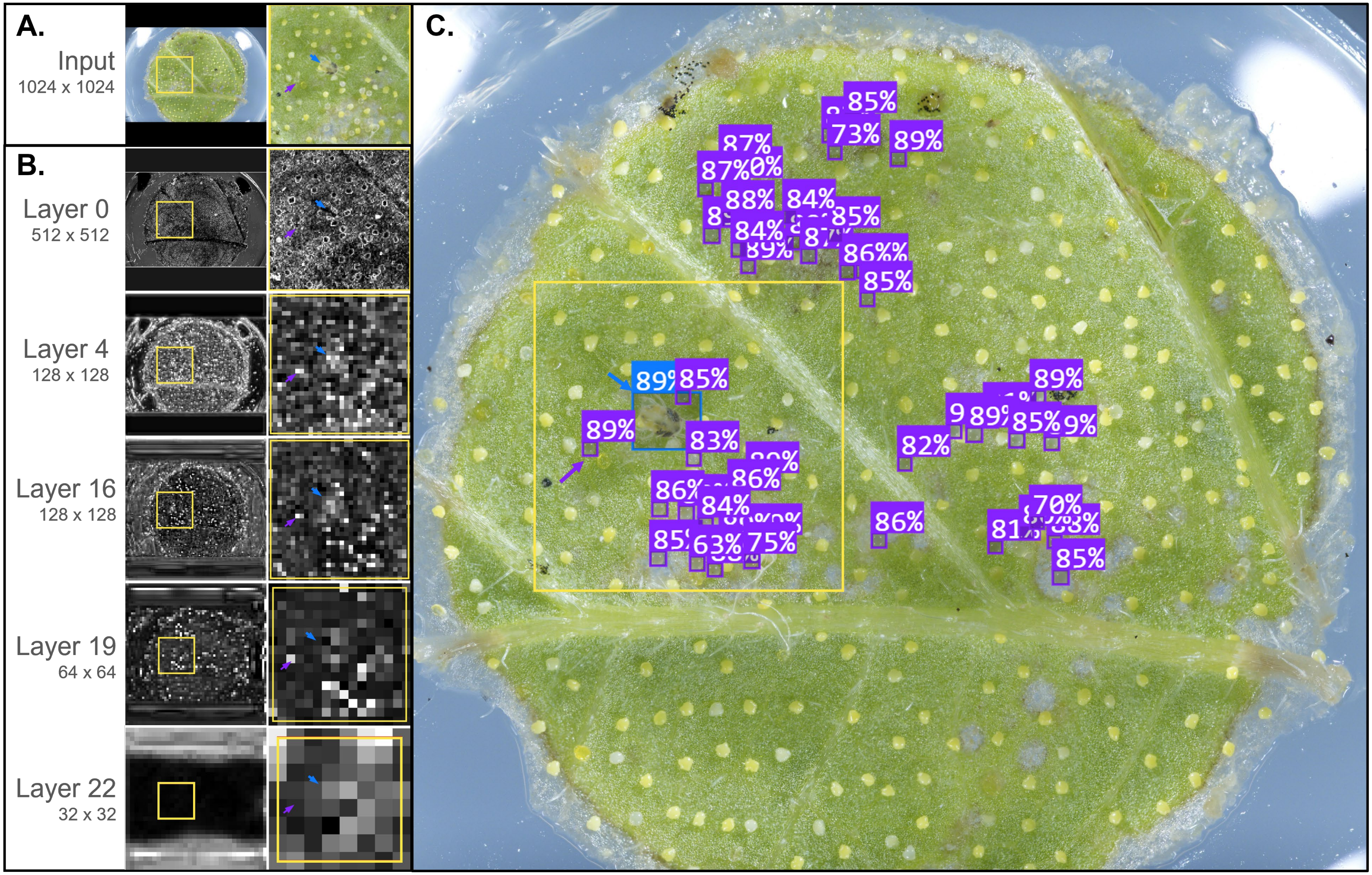

**S1 Fig. Hierarchical feature representations and detection outputs from the YOLO11-L model.**
**(A)** Input image with the region of interest highlighted (yellow box) for visualization of internal feature activations.
**(B)** Representative channel-averaged feature maps from selected stages of the YOLO11 architecture. Early backbone layers (e.g., Layer 0) show higher-resolution responses to local texture and edges, whereas deeper backbone layers (e.g., Layer 4 to 8) exhibit progressively more abstract, semantically integrated representations. Neck layers and detection-head input scales (Layers 16, 19, and 22) correspond to the three feature-pyramid levels used for object detection (P3, P4, and P5, respectively), with P3 providing the highest spatial resolution used for detecting very small objects. For each layer, the right-hand inset shows the same region of interest at the native feature-map resolution. Blue and purple arrows indicate an *Adult_female* (blue) and a *Viable_egg* (purple). Areas of lighter intensity represent stronger activation responses. **(C)** Final detection output for the same image, showing bounding boxes and confidence scores for *Adult_female* (blue) and *Viable_egg* (purple) detections.

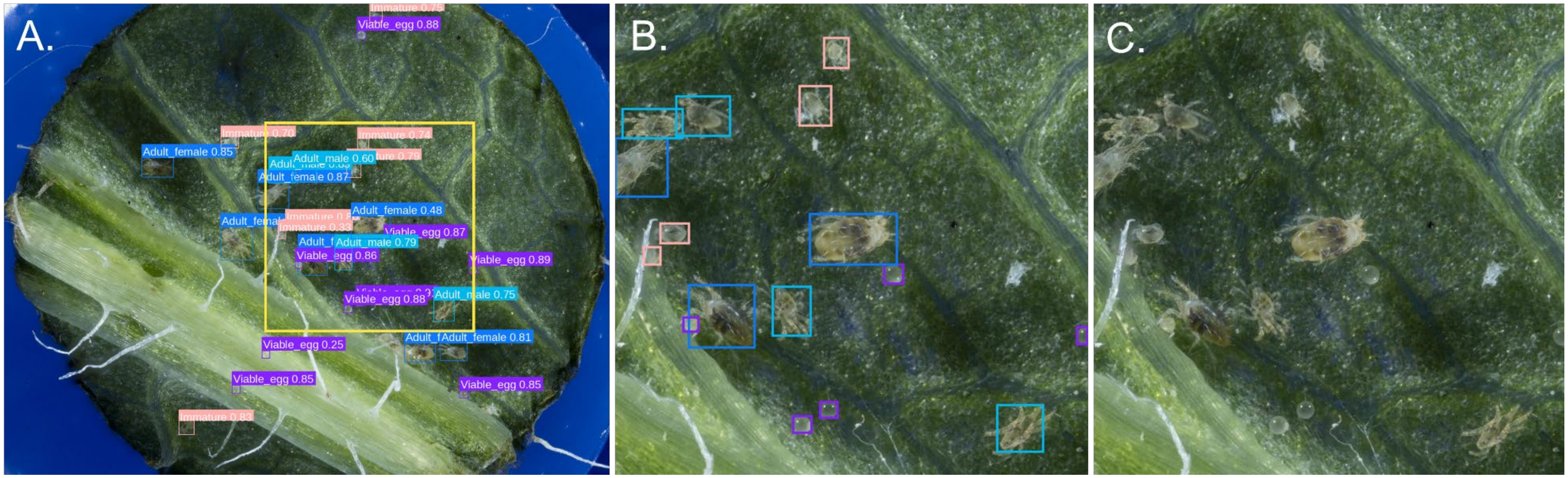


**S2 Fig. Example performance of the four-class detection model (v211) on a test-set image.**
**(A)** Model inference results (IoU = 0.5, confidence threshold = 0.5) showing detected objects with labels and confidence scores. Detected classes are color-coded as follows: *Adult_female* (blue), *Adult_male* (teal), *Immature* (salmon), and *Viable_egg* (purple). **(B)** Magnified view of the region outlined in panel A, illustrating the model's ability to detect partially occluded and moving individuals. **(C)** The same magnified region shown without labels for visual comparison.

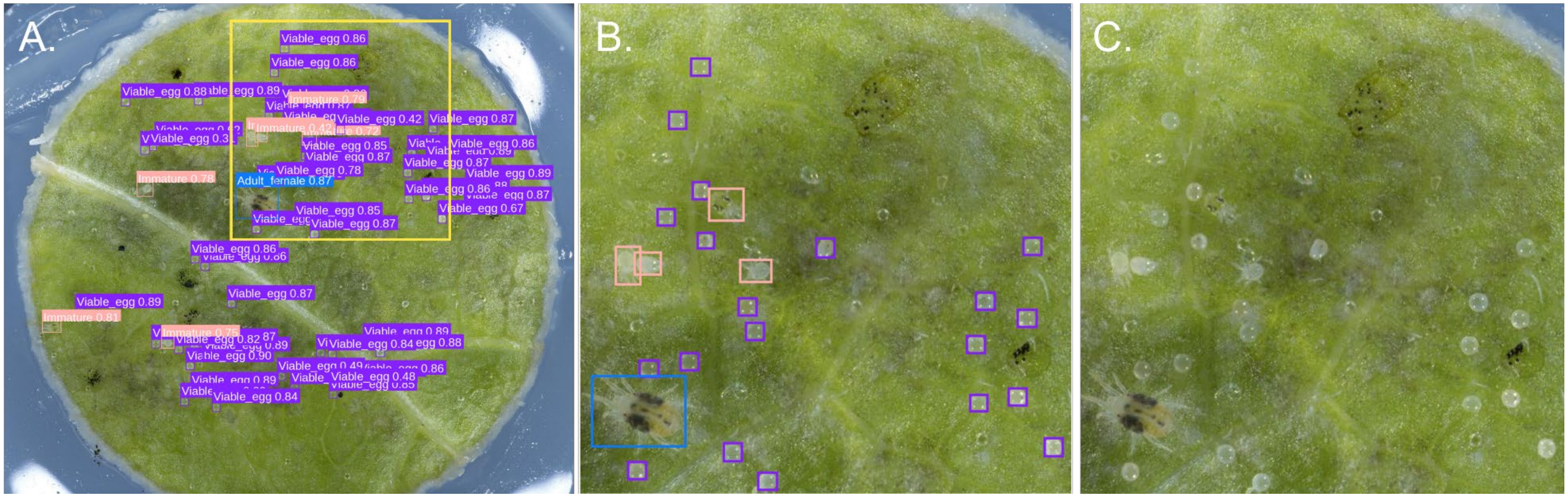


**S3 Fig. Example performance of the three-class detection model (v210) on a test-set image.**
**(A)** Model inference results (IoU = 0.5, confidence threshold = 0.5) showing detected objects with labels and confidence scores. Detected classes are color-coded as follows: *Adult_female* (blue), *Immature* (salmon), and *Viable_egg* (purple). **(B)** Magnified view of the region outlined in panel A. **(C)** The same magnified region shown without labels for visual comparison.

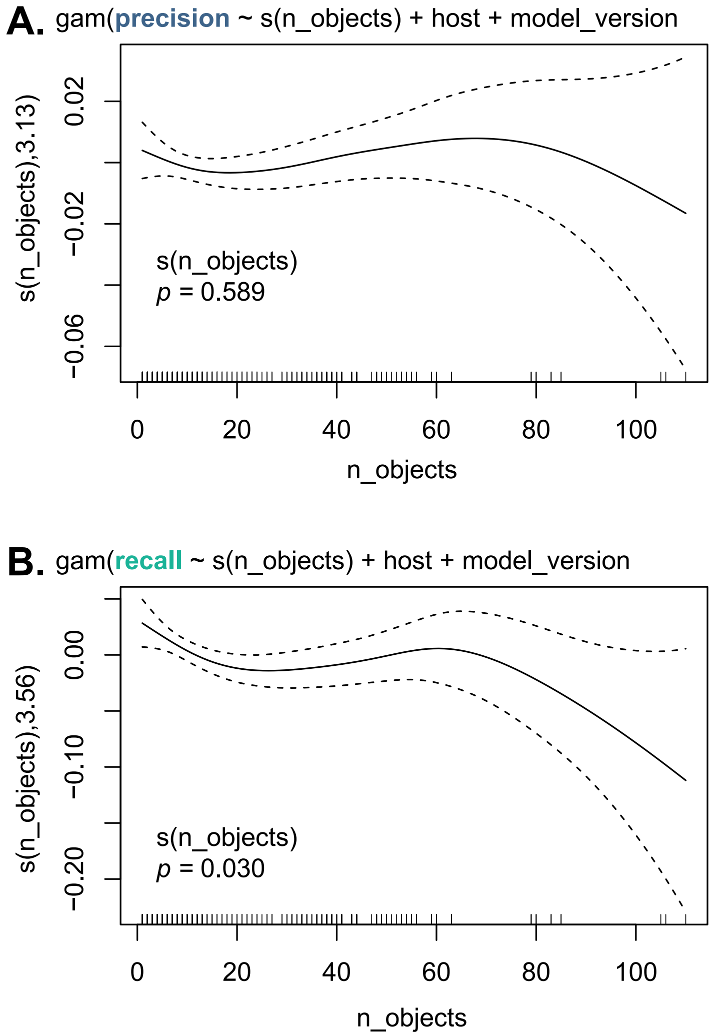


**S4 Fig. Generalized additive model smooth terms for the effect of object count on per-image precision and recall. (A)** Precision showed no significant association with object count ($p = 0.589$), indicating stability across varying object densities. **(B)** Recall exhibited a significant nonlinear relationship with object count ($p = 0.030$), with performance declining at higher object counts. Dashed lines indicate approximately 95% confidence intervals (±2 standard errors).

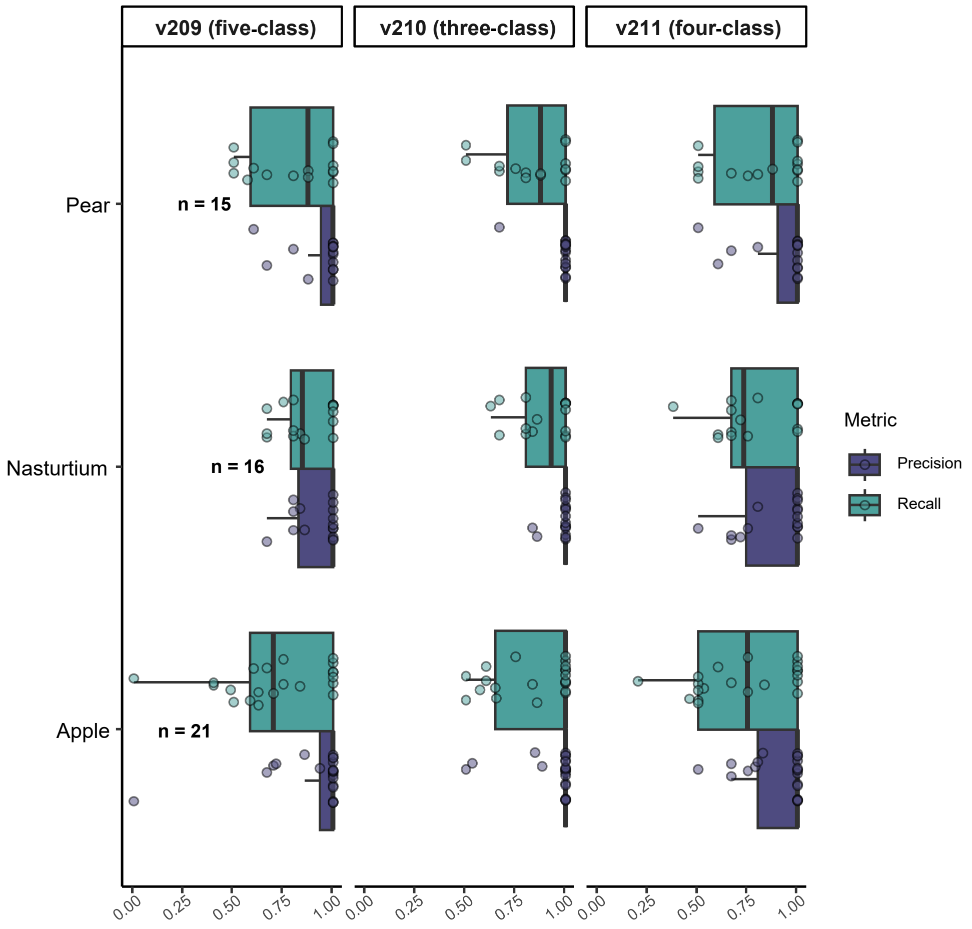


**S5 Fig. Per-image precision and recall for hosts not included in model training.** Performance metrics are shown for three previously unseen host plants (apple, nasturtium, and pear) across all three model versions (209, 210, and 211). Each point represents a single image, with boxplots summarizing distributions of precision and recall per host. While precision remains consistently high across hosts, recall shows greater variability, with apple exhibiting the lowest recall values overall. Sample sizes (n) are indicated next to each group.

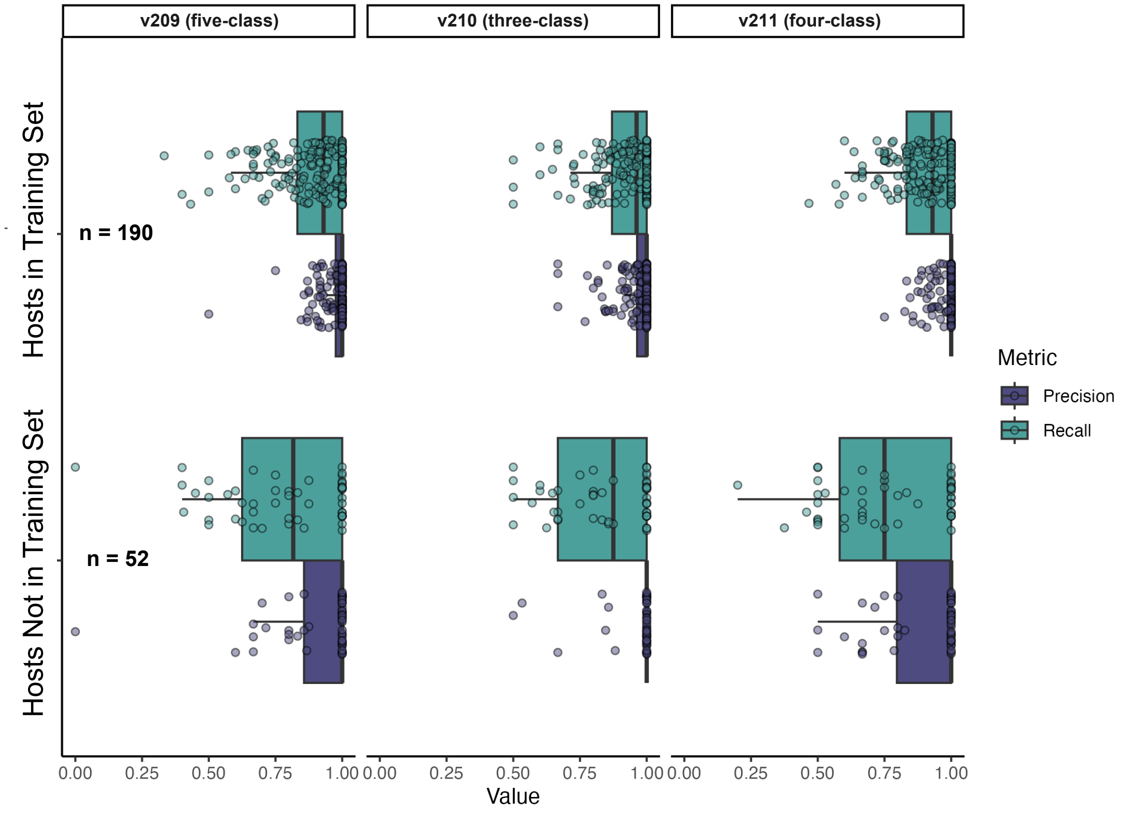


**S6 Fig. Performance of three model versions (v209: five-class; v210: three-class; v211: four-class) on the leaf test set, comparing hosts included in training ("pretrained") versus previously unseen hosts.** Each point represents a test image, with precision (purple) and recall (green) shown. Recall dropped markedly for unseen hosts, whereas precision was largely unaffected, indicating greater sensitivity of recall to domain shift. Sample sizes (n) are shown for each group. Training hosts: hop, tomato, bean, raspberry, columbine, poppy, strawberry; unseen hosts: apple, pear, nasturtium.

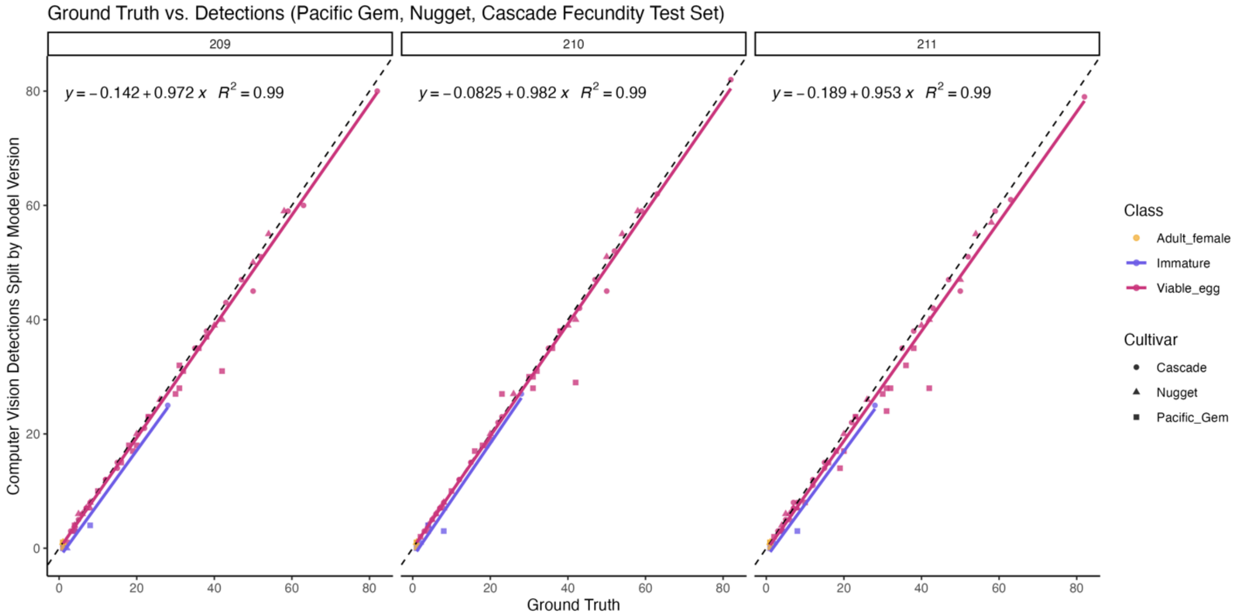


**S7 Fig. Predicted vs. ground truth mite detections on a subset of hop mite fecundity study.** Predicted object counts from three computer vision models are plotted against ground truth data from a subset of the cultivar fecundity study, encompassing three hop cultivars (Cascade, Nugget, and Pacific Gem). Points are colored by mite class and faceted by model version and cultivar. The dashed diagonal black line represents 1:1 agreement; solid lines denote linear regression fits. The regression equation and $R^2$ shown in each panel represent a single overall linear fit computed from all points in that facet, pooled across mite classes and cultivars.

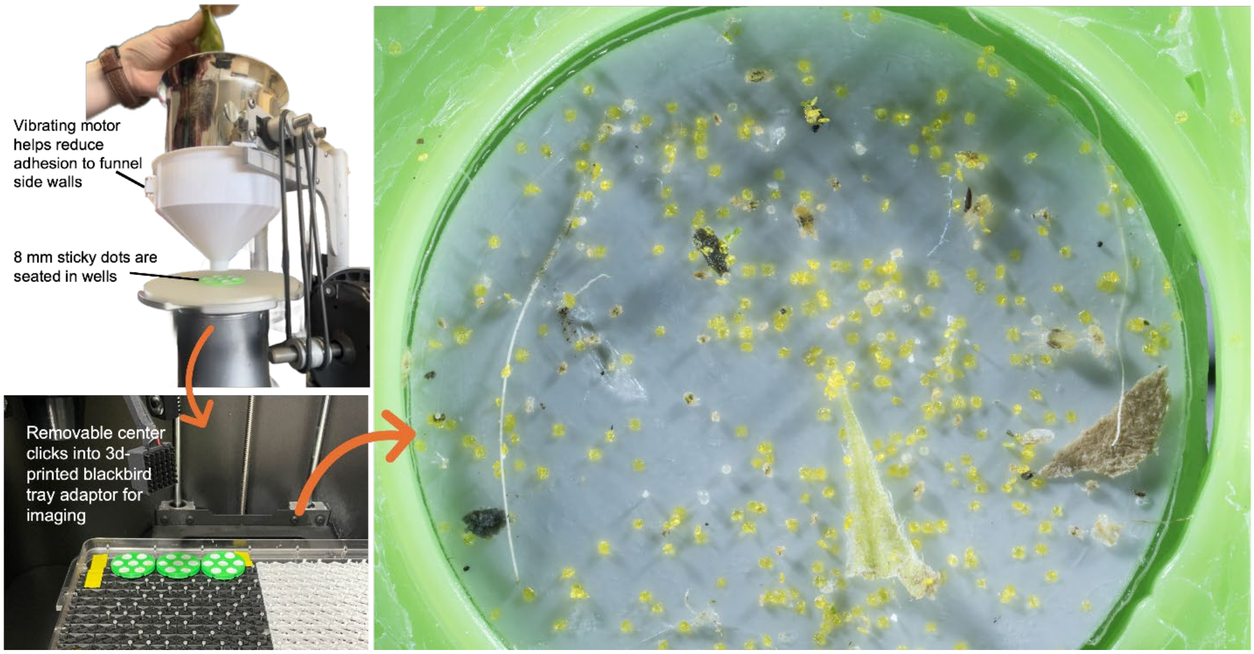


**S8 Fig. Adapting mite imaging pipeline for field-collected mites.** We concentrated mite brushing products for imaging with the Blackbird platform by 3D printing custom-fit funnels (upper left) for the mite brushing machine and 8 mm sticky dot sample trays (lower left). Unfortunately, we found that mite brushing often results in mite fatalities, complicating efforts to infer their on-leaf viability for both human observers and computer vision models. Additionally, debris from the leaf samples often occluded target objects further reducing accuracy and recall performance (as seen on the right).

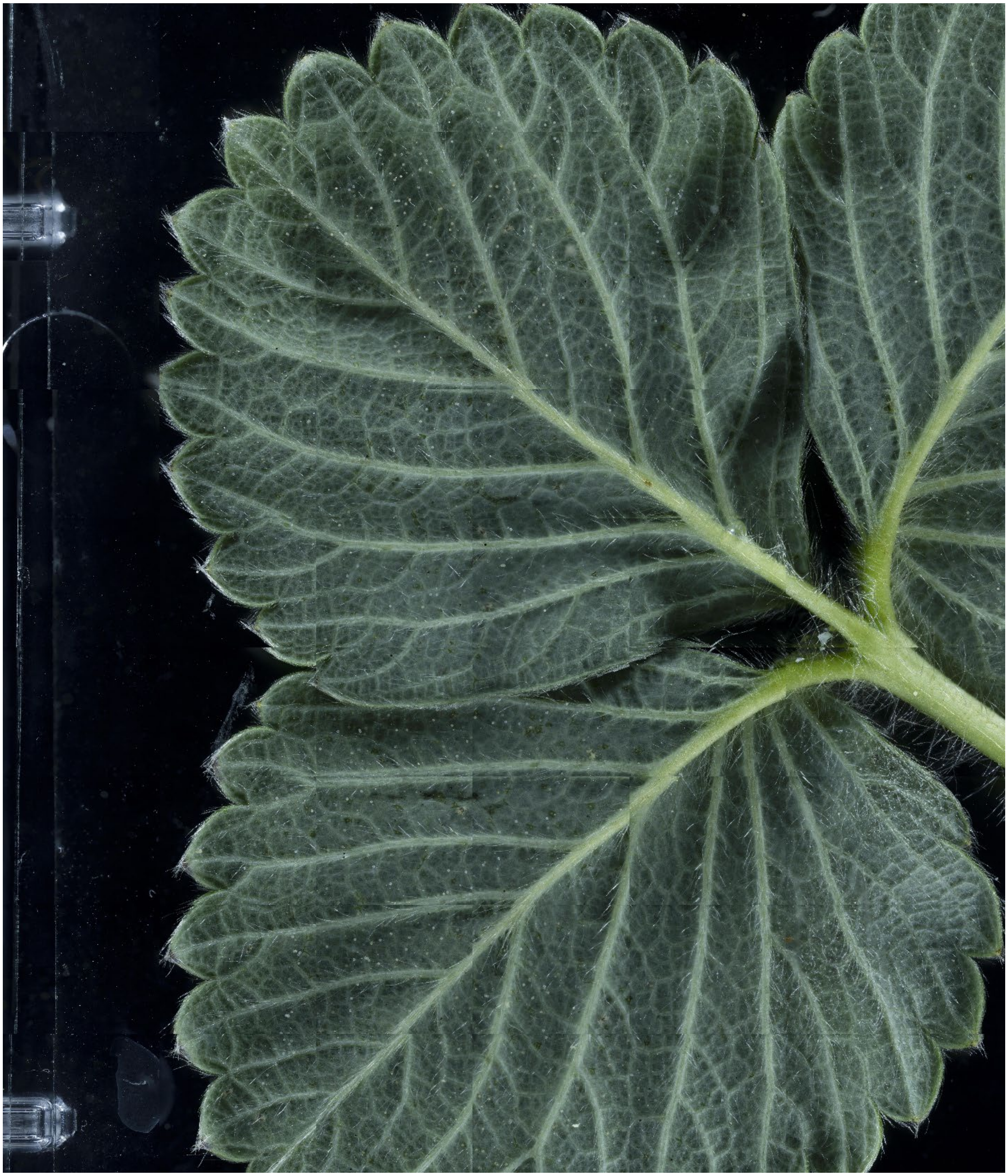

**S9 Fig. Leaf-scanning for computer vision detection on field-derived material.** To address challenges posed by occlusion and debris following mite brushing and concentration, we used the Blackbird imaging system to tile full leaf surfaces in a "scanning" mode. This image of a field-collected strawberry leaf comprises 54 stitched z-stacks, totaling approximately 3.78 GB of raw image data. Although this method improves detection accuracy on complex, field-derived samples, it is not practical for large-scale use due to the time required and the extremely large file sizes.

### Supplemental tables

**Table S1. Performance comparison of YOLO variants, RT-DETR-L, and Faster R-CNN with two backbones on the v209 training and test sets.** Highest performance in each column is indicated in bold.

| Rank | Model | Framework | mAP50-95 | mAP50 | FPS |
|---|---|---|---|---|---|
| **1** | **YOLO11-L** | Ultralytics | **0.535** | **0.836** | 50.9 |
| 2 | YOLOv8-L | Ultralytics | 0.510 | 0.830 | 67.4 |
| 3 | RT-DETR-L | Ultralytics | 0.486 | 0.770 | 36.7 |
| 4 | YOLOv8-N | Ultralytics | 0.454 | 0.752 | **109.8** |
| 5 | YOLO11-N | Ultralytics | 0.435 | 0.743 | 80.6 |
| 6 | FasterRCNN-R50-FPNv2 | Torchvision | 0.336 | 0.588 | 42.8 |
| 7 | FasterRCNN-MobileNetV3-FPN | Torchvision | 0.098 | 0.244 | 58.4 |

**Table S2. Cross-validation fold results for the five-class model (v209).** Performance metrics averaged across all classes.

| Fold | Precision | Recall | mAP50 | mAP50-95 |
|---|---|---|---|---|
| 0 | 0.824 | 0.761 | 0.824 | 0.528 |
| 1 | 0.849 | 0.703 | 0.793 | 0.488 |
| 2 | 0.808 | 0.711 | 0.781 | 0.457 |
| 3 | 0.758 | 0.756 | 0.783 | 0.495 |
| 4 | 0.809 | 0.815 | 0.855 | 0.528 |
| Mean | 0.810 | 0.749 | 0.807 | 0.499 |

**Table S3. Mean per-class performance metrics across all *k*-folds for the five-class model (v209).**

| Class | Precision | Recall | mAP50 | mAP50-95 |
|---|---|---|---|---|
| Adult_female | 0.864 | 0.893 | 0.927 | 0.585 |
| Adult_male | 0.675 | 0.682 | 0.720 | 0.401 |
| Dead_mite | 0.649 | 0.460 | 0.540 | 0.316 |
| Immature | 0.898 | 0.788 | 0.877 | 0.490 |
| Viable_egg | 0.961 | 0.924 | 0.972 | 0.706 |

**Table S4. Resolution limit testing aggregate results.** Highest performance in each column is indicated in bold.

| Native Resolution | Precision | Recall | mAP50 | mAP50-95 |
|---|---|---|---|---|
| 1024 × 1024 (100%) | 0.888 | **0.765** | **0.840** | **0.577** |
| 819 × 819 (80%) | **0.892** | 0.715 | 0.814 | 0.562 |
| 512 × 512 (50%) | 0.866 | 0.555 | 0.719 | 0.476 |
| 256 × 256 (25%) | 0.694 | 0.241 | 0.461 | 0.251 |

**Table S5. mAP50 resolution limit testing results by class.** Highest performance in each column is indicated in bold.

| Native Resolution | Adult_female | Adult_male | Dead_mite | Immature | Viable_egg |
|---|---|---|---|---|---|
| 1024 × 1024 (100%) | **0.912** | **0.806** | **0.674** | **0.881** | **0.928** |
| 819 × 819 (80%) | 0.891 | 0.783 | 0.624 | 0.873 | 0.902 |
| 512 × 512 (50%) | 0.855 | 0.679 | 0.562 | 0.766 | 0.734 |
| 256 × 256 (25%) | 0.749 | 0.372 | 0.330 | 0.401 | 0.454 |
